# Herbivory enhances leaky sex expression in the dioecious herb *Mercurialis annua*

**DOI:** 10.1101/2021.04.19.440496

**Authors:** Nora Villamil, Xinji Li, Emily Seddon, John R. Pannell

## Abstract

Plant reproductive traits are widely understood to be responsive to the selective pressures exerted by pollinators, but there is also increasing evidence for an important role of antagonists such as herbivores in shaping these traits. Many dioecious species show leaky sex expression, with males and females occasionally producing flowers of the opposite sex. Here we show that leaky sex expression in both males and females of the wind-pollinated dioecious herb *Mercurialis annua* (Euphorbiaceae) is enhanced in response to simulated herbivory, increasing the probability and the degree of leakiness in both sexes. We also found that leakiness was greater in larger females but not in larger males. We discuss hypotheses for a possible functional link between herbivory and leaky sex expression, and consider what herbivory-induced leakiness might imply for the evolutionary ecology of plant reproductive systems, especially the breakdown of dioecy and the evolution of hermaphroditism.

## Introduction

Floral morphology and other reproductive traits are often strongly influenced by selection due to interactions with pollinators (Willmer 2011). However, reproductive traits can also be shaped by selection due to antagonists such as herbivores (Ashman 2002, Steets and Ashman 2004, Carr and Eubanks 2014, Johnson et al. 2015). For instance, the strength of selection due to herbivores was found to be equal to or stronger than that due to pollinators in 67% of species in which the role of both mutualists and antagonists had been studied (Johnson et al. 2015). There is also growing evidence for the coordinated evolution of plant defensive and reproductive traits (Strauss and Whittall 2006, Rausher 2008, Ågren et al. 2013, Campbell and Kessler 2013, Carr and Eubanks 2014, Campbell 2015, Johnson et al. 2015). Specifically, herbivores, and plant responses to them, have been found to affect the floral display (Strauss et al. 1996, Strauss et al. 1999, Herrera 2000, Parachnowitsch and Caruso 2008, Ågren et al. 2013), flower colour (Strauss et al. 2004, Vaidya et al. 2018), flowering time (Parachnowitsch and Caruso 2008), floral morphology (Galen 1999, Galen and Cuba 2001, Santangelo et al. 2019), floral scents and pollinator rewards (Kessler and Halitschke 2009, Kessler et al. 2011, Ramos and Schiestl 2019, Aguirre et al. 2020), mating systems (Steets and Ashman 2004, Ivey and Carr 2005, Penet et al. 2009, Kessler et al. 2011), and sex allocation (reviewed by Ashman 2002). Yet despite the increasing evidence for the importance of herbivory in the ecology and evolution of plant reproduction, very little attention has been given to the effect of herbivory on sex allocation and other traits that might be important in evolutionary transitions between hermaphroditism and dioecy.

Herbivores can affect sex allocation patterns either directly via plant size or indirectly via effects on pollinator behaviours. By reducing plant size and resource availability, herbivory in dioecious populations may directly increase mortality of individuals with the more costly sex function, potentially leading to or enhancing sex ratio biases (Cornelissen and Stiling 2005, Sánchez-Vilas and Pannell 2011, Geber et al. 2012). For similar reasons, herbivory in hermaphroditic species can cause individuals to shift their investment towards the least costly sexual function (Diggle 1994, Seger and Eckhart 1996, Ashman 2002, Zhang and Jiang 2002, West 2009, Hirata et al. 2019). A particularly interesting and important possibility is that herbivory could influence the sex allocation of males and females of dioecious species by altering their tendency to produce flowers of the opposite sex through ‘leaky’ or ‘inconstant’ sex expression.

Leaky sex expression has been described in at least 40 dioecious species, including gymnosperms and angiosperms (Ehlers and Bataillon 2007, Cossard and Pannell 2019), but it is probably more frequent. In some cases, it may simply reflect the rudimentary expression of hermaphroditism in species that have not completed the transition to fully separate sexes (Delph 2003; Delph and Wolf 2005). However, it may also be maintained by selection for reproductive assurance under conditions in which mating opportunities are limited (Crossman and Charlesworth 2014, Cossard et al. 2021, Cossard and Pannell 2021). Mate limitation could arise from colonising new, mate-less environments (Baker 1965, Pannell 2015) or from pollinator limitation caused by biotic (e.g. herbivory) or abiotic factors (e.g. environmental heterogeneity, such as across altitudinal gradients) (Kessler et al. 2011, Trøjelsgaard and Olesen 2013). Until recently, almost nothing was known about the developmental basis of leakiness, but several studies indicate that it may have a plastic component, with plants responding to biotic or abiotic factors (Freeman et al. 1980, Bierzychudek and Eckhart 1988, Sakai et al. 1995). For example, in some species cool and humid conditions at high altitudes have been found to favour males and strict dioecy (unisexuality), whereas warmer, drier conditions at lower altitudes favour hermaphrodites and enhanced leakiness, showing leakiness response to environmental cues (Delph 1990b, Sakai and Weller 1991, Humeau et al. 1999, Venkatasamy et al. 2007). A recent study of leaky sex expression in the wind-pollinated dioecious plant *Mercurialis annua* found that leakiness can also respond plastically to plant-plant interactions and population sex ratios (Cossard and Pannell 2021), such that females deprived of pollen-producing mates are more likely to produce male flowers than comparable females receiving abundant pollen (Cossard and Pannell 2021).

Here, we asked to what extent leakiness in sex expression in *M. annua* might also be plastically responsive to herbivory. This is important because enhanced leakiness in dioecious populations could lead to a shift in both the mating system and in the conditions for transitions between combined and separate sexes. Our study also addresses a perceived gap in our understanding, identified by Johnson et al. (2015), of how herbivore-induced changes in phenotype might alter a species’ mating system. Most documented cases of the effect of herbivory on the mating system point to reduced selfing in hermaphrodite species (reviewed by Johnson et al. 2015), whereas enhanced leakiness in dioecious species would potentially allow selfing instead. Previously, Yampolsky (1930) and Kuhn (1939) found that pruning individuals of *M. annua* tended to enhance the production of flowers of the opposite sex, but neither study characterised this response in any detail.

Herbivory might affect leakiness in sex expression for a number of reasons. First, a plant’s response to herbivory might include altering its endogenous hormone balance (Thaler et al. 2001, Ballaré 2011, Robert-Seilaniantz et al. 2011, Naseem et al. 2015), with potential pleiotropic effects on its sex expression (Riemann et al. 2003, Wasternack et al. 2013, Yuan and Zhang 2015). And second, herbivory might affect a plant’s sex expression via effects on plant size or resource status, as predicted by theories of size- or resource-dependent sex allocation (Ghiselin 1969, Trivers and Willard 1973, Charnov 1979, Freeman et al. 1980, Charnov 1982, Warner 1988, de Jong and Klinkhamer 1989, Klinkhamer et al. 1997, reviewed by West 2009). For instance, if physical injuries such as those caused by herbivory reduce an individual’s resource status, selection might favour a strategy that includes a shift in sex allocation towards the cheaper sex function. While studies of size- and/or resource-dependent sex allocation and gender (Lloyd 1980) have tended to focus on hermaphrodites (Freeman et al. 1980, Korpelainen 1998, VegalJFrutis et al. 2014), the same ideas might apply to sex inconstancy in dioecious species.

We examined the effect of simulated herbivory on the sexual expression of males and females of *M. annua* in two experiments in which different levels of simulated herbivory led to enhanced leakiness in both sexes. In females, we compared leakiness levels between control (undamaged) and damaged females. In males, we quantified leakiness of plants under a low and a high herbivory treatment. We used the data from both experiments to address the following questions: (1) How does herbivory affect the probability of leakiness in males and females of a dioecious species? (2) How does herbivory affect the number of opposite sex reproductive structures produced by males and females of a dioecious species? (3) To what extent are the male and female changes in sex expression in response to herbivory mediated by plant size?

## Materials and methods

### Study system

*Mercurialis annua* is a polyploid complex of wind-pollinated ruderal herbs that occupy disturbed habitats across eastern, central and western Europe (Tutin et al. 1968, Obbard et al. 2006). Diploid populations are dioecious, with an XY chromosomal system of sex determination (Russell and Pannell 2015, Veltsos et al. 2018, Li et al. 2019, Veltsos et al. 2019). Males produce staminate flowers on pedunculate inflorescences held above the plant. Females produce two-to three-ovulate flowers on sub-sessile pedicels in the leaf axils (Tutin et al. 1968). Apart from these differences in the floral sex and inflorescence morphology between males and females, the sexes also differ in a number of vegetative characters, including plant and root biomass, patterns of resource allocation to growth and reproduction throughout their development, and their competitive abilities (Sánchez□Vilas et al. 2011, Sánchez Vilas and Pannell 2011, Tonnabel et al. 2017). As in the case of many dioecious plants (Ehlers and Bataillon 2007, Cossard and Pannell 2019), dioecious *M. annua* shows leakiness in sex expression, with both males and females occasionally producing fully functional flowers of the opposite sexual function (Pannell et al. 2008, Cossard and Pannell 2019, 2021).

### Plant culture

Plants of the diploid *M. annua* were sown and grown within a polytunnel under controlled conditions at the University of Lausanne, Switzerland. The experiment with female plants was established during March 2016, while the experiment with male plants was established in May 2016. In both experiments, plants were sown and raised to maturity in seedling trays. When plants reached reproductive maturity, plants of the desired sex for the respective experiment were repotted in pots with soil (Ricoter substrate 140) and slow-release fertilizer (Hauert Tardit 6M pellets; 5 g fertilizer/L of soil). Plants were subjected to herbivory treatments and allowed to regrow for 10 weeks in the male experiment and 8 weeks in the female experiment. After this period, plants were harvested, and the numbers of male and female flowers were recorded. Plants were then dried and weighed.

### Herbivory treatments

For the male experiment, individuals were grown in pairs in pots to save space (with pot thus being the unit of replication; n = 828 pots). Both individuals in a given pot were subjected to the same low- or high-herbivory treatment. Over a period of 10 weeks of growth, plants under the low-herbivory treatment were pruned once at the first internode (3 cm above the soil surface), whereas plants under the high-herbivory treatment were pruned twice at the first internode. For the female experiment, individuals were transplanted into individual pots (n = 219 plants). For plants under the herbivory treatment, the apical section (10 cm) of the main stem was pruned once, whereas plants under the control treatment were left intact.

Our experiment on male plants involved a comparison between two treatments of simulated herbivory of contrasting intensity, but did not include undamaged plants. This is because it was initially part of a study that specifically aimed at generating males with leaky sex expression for the production of YY male plants (Li et al. 2019). The absence of undamaged males means that our low *vs*. high herbivory comparison is a more conservative estimate of sensitivity to the intensity of herbivory, as confirmed by previously documented levels of leaky sex expression for undamaged males of the same population (Cossard and Pannell 2019, see Discussion for details). The male experiment thus specifically asks how sensitive plants are in their leakiness to the degree of herbivory rather than to herbivory as a categorical variable.

### Statistical analyses

Statistical analyses were conducted in R version 3.6.1 (R Core Team 2016). Models were fitted using ‘lme4’ R package (Bates et al. 2016), unless otherwise stated, with residuals evaluated with the ‘DHARMa’ R package (Hartig and Hartig 2017).

To test whether the herbivory treatment affected the probability of sex change in males and females, we fitted generalised linear binomial models. For males, the presence or absence of seeds was considered as the response variable, with the herbivory treatment and biomass fitted as predictors. For females, the presence or absence of male flowers was the response variable, and the predictors were the herbivory treatment, plant biomass, and plant population.

To test the effect of herbivory on the number of reproductive structures of the opposite sex, we fitted two separate Poisson models. We tested and accounted for zero-inflation in both models. Seed production in males was zero-inflated and was consequently analysed with a Poisson zero-inflated mixed model using the ‘glmmADMB’ package (Skaug et al. 2011). The number of seeds produced by males was fitted as the response variable, with the herbivory treatment and plant biomass fitted as fixed effects. To control for overdispersion, we also included an observation-level random effect (OLRE) (Hinde 1982, Harrison 2014). The model testing the effect of herbivory on male flower production by females was not zero-inflated. Here, we fitted the number of male flowers produced by females as the response variable, the herbivory treatment and plant biomass as fixed effects, and included an observation-level random effect (OLRE).

## Results and Discussion

### Patterns of leakiness in sex expression in M. annua

Herbivory significantly increased the probability and the degree of leakiness in both males and females (Tables 1, S1, Fig. 1). Thus, males in pots under high herbivory were 15% more likely to produce seeds than those under low herbivory (Fig. 1A) and they produced 13 times more seeds on average (Fig. 1C, Table 1). Similarly, females under herbivory were 26% more likely to produce male flowers than control females (Fig. 1B), and they produced five times more male flowers than control females (Fig. 1D, Table 1). We also found that while females were 0.51% more likely to produce male flowers and produced 1.06 more male flowers per gram of additional biomass, male leakiness did not depend on plant size (Table 1).

**Table 1.**
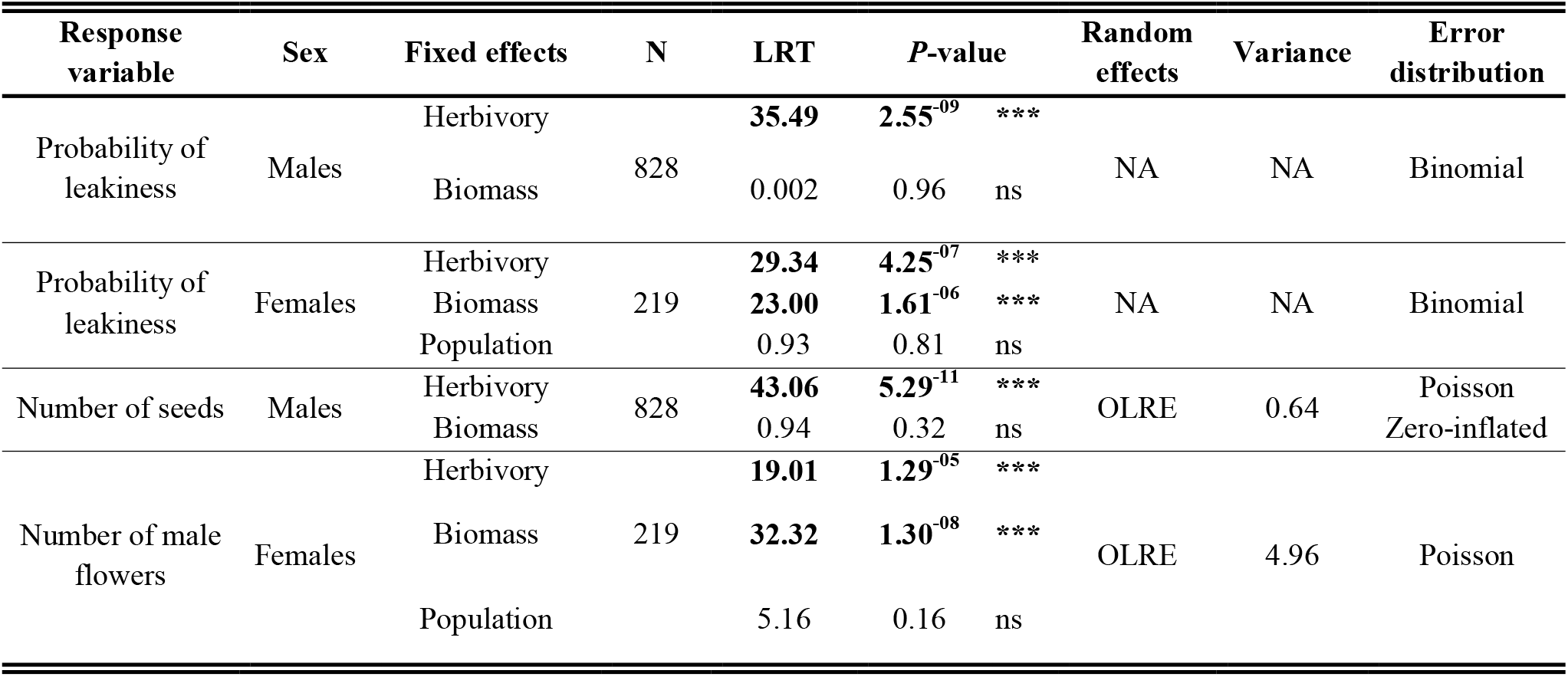
Model output for the effects of simulated herbivory on leakiness in sex expression in males and females of *Mercurialis annua*. LRT: Likelihood ration test; OLRE: observation-level random effect, added to control for overdispersion in Poisson models.

**Table S1.** Estimates based on model predictions: indicate are the mean, standard error and the lower and upper 95% confidence intervals.

| Response variable | Sex | Treatment | Mean | SE | Lwr 95 CI | Upr 95 CI |
| --- | --- | --- | --- | --- | --- | --- |
| **Leakiness probability** | Males | Low herbivory | 0.045 | 0.25 | 0.028 | 0.072 |
|  |  | High herbivory | 0.193 | 0.134 | 0.155 | 0.237 |
|  | Females | Control | 0.351 | 0.216 | 0.261 | 0.453 |
|  |  | Herbivory | 0.614 | 0.207 | 0.514 | 0.705 |
| **Number of seeds** | Males | Low herbivory | 0.241 | 0.562 | 0.080 | 0.723 |
|  |  | High herbivory | 1.806 | 0.312 | 0.981 | 3.325 |
| **Number of male flowers** | Females | Control | 0.271 | 0.324 | 0.143 | 0.513 |
|  |  | Herbivory | 1.366 | 0.262 | 0.816 | 2.287 |

**Figure 1.**
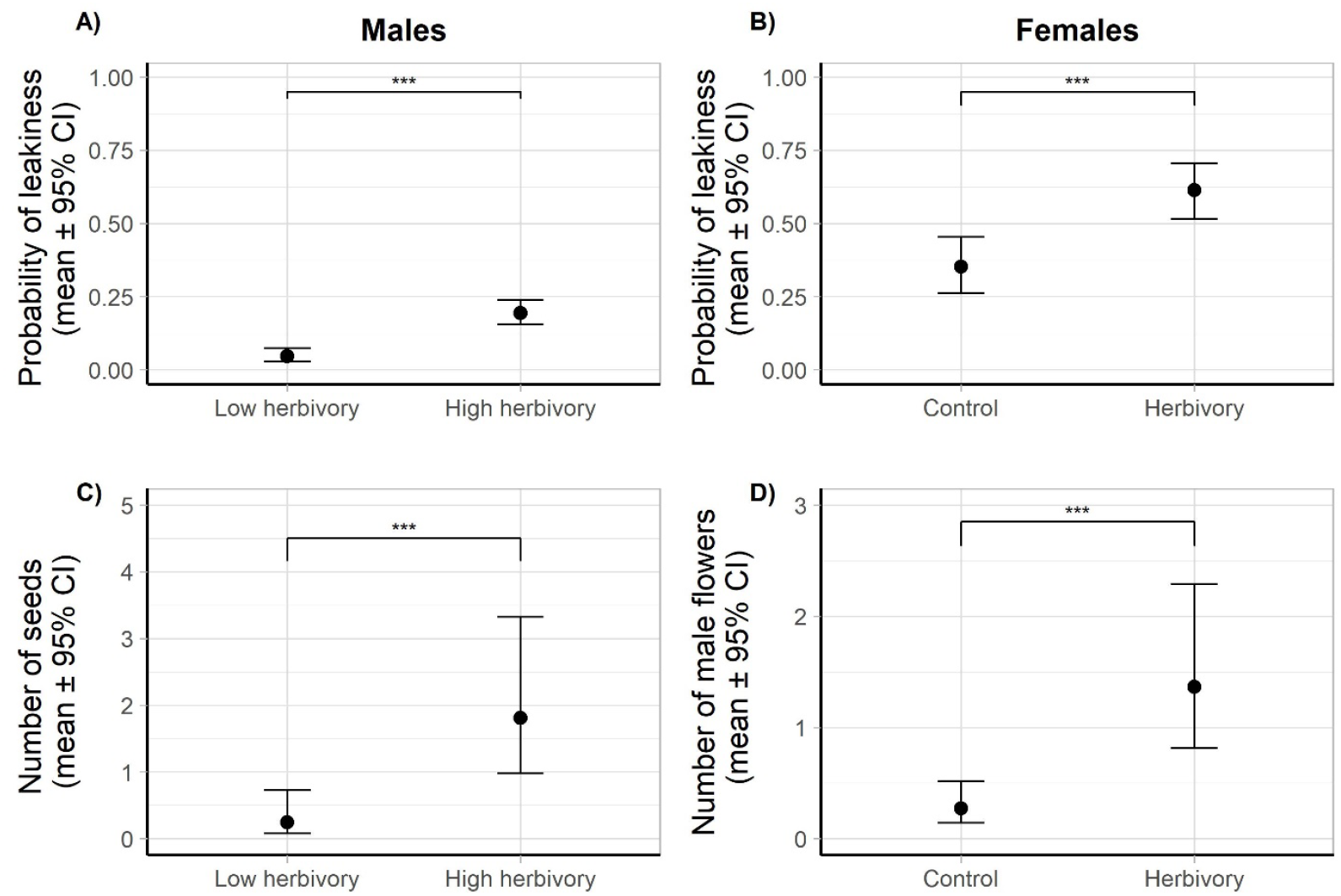
Leakiness in sex expression in response to simulated herbivory for males and females of *Mercurialis annua*, in terms of effects on the probability of leakiness (A-B), and in the number of reproductive structures of the opposite sex (C-D). Error bars represent 95% confidence intervals and asterisks indicate significant differences (P < 0.0001).

While leakiness in sex expression can affect both sexes of dioecious species, generally males are more likely to be leaky than females, a pattern that may reflect incomplete transitions from hermaphroditism to dioecy via gynodioecy, with males retaining a residual female function (Ehlers and Bataillon 2007, Cossard and Pannell 2019). Against this background, the greater probability of leakiness in females of *M. annua* is unusual. Our results confirm greater leakiness in females than males in *M. annua*, and they also indicate that under herbivory this pattern of enhanced leakiness is maintained and even accentuated. Previous data on the baseline probability of sex inconstancy in undamaged *M. annua* males of the same source population were 3% (Cossard and Pannell 2019), compared with 4.5% in males under the low herbivory and 23.8% under the high herbivory treatment.

Dioecy is ancestral and well-established across the genus *Mercurialis*, and there is no indication that separate sexes evolved via gynodioecy (though they might have done so). Greater leakiness in females than males might reflect selection for reproductive assurance, which would favour the maintenance of leaky sex expression in females rather than males, because a small amount of pollen produced by females might suffice for the production of a large number of seeds, in contrast with the few seeds produced by males (Pannell and Barrett 2001, Cossard and Pannell 2019). This might also explain why we found that males, when leaky, tend to express a greater degree of leakiness than females, as found previously by Cossard and Pannell (2019). These patterns of leakiness in *M. annua* are thus consistent both with the expectations of range-expansion or metapopulation models, in which new populations are frequently established by single individuals, as well as with the ruderal habit and metapopulation structure of *M. annua* across Europe (Pannell and Barrett 2001, Obbard et al. 2006, Eppley and Pannell 2007b).

Our results shed further light on the expression of leakiness in sex expression in *M. annua*, to our knowledge the only plant to date whose leakiness has been investigated experimentally in quantitative terms, showing that antagonists can affect this reproductive trait. Leaky sex expression is common in dioecious species, but the basis of variation in leakiness among individuals has remained almost entirely obscure. Our current and previous studies of leaky sex expression in *M. annua* indicate that the phenomenon cannot be attributed only to developmental instability or poorly canalised separation of the sexes. Rather, leakiness is clearly a more complex trait with components of variation attributable to not only genetic differences among individuals (Cossard et al. 2021), but also to phenotypic plasticity. Previous work found that females of *M. annua* were more likely to express a male function when growing under conditions of pollen (or mate) limitation (Cossard and Pannell 2021). By showing that leaky sex expression in *M. annua* also responds to herbivory, our current study now adds further evidence for the contribution of phenotypic plasticity to phenotypic variation in sex expression and highlights the role antagonists can have in shaping reproductive traits in angiosperms. Our findings also provide evidence reinforcing the idea of coordinated evolution between defensive and reproductive traits in angiosperms showing herbivory-induced leakiness in male and female plants.

### Why should leakiness in sex expression be sensitive to herbivory?

While it seems plausible that the sensitivity of leaky sex expression of unisexual plants to mate limitation might have evolved in response to selection for reproductive assurance in *M. annua* (Cossard and Pannell 2021), a functional explanation for the sensitivity of leakiness to simulated herbivory is less obvious. Could it be that herbivory-enhanced leakiness functions as a reproductive assurance mechanism too, e.g., if herbivory compromised mate availability? In most dioecious species, herbivory is male-biased (Ågren et al. 1999, Cornelissen and Stiling 2005, Geber et al. 2012), including in *M. annua* (Sánchez-Vilas and Pannell 2011), so that at least females might gain from induced leakiness in strongly damaged populations. Hesse and Pannell (2011) found that isolated females of *M. annua* were indeed pollen limited in the field, but we have no evidence that herbivory frequently brings about such isolation in *M. annua*, nor indeed how effectively the observed levels of leakiness in our experiment would actually restore seed production to pollen-limited females.

Another possibility is that enhanced leakiness in response to herbivory might be a collateral effect of hormonal changes resulting from the activation of plant defensive pathways. Plant hormones are known to regulate both sex determination and defensive responses in a number of plants (Robert-Seilaniantz et al. 2011, Yuan and Zhang 2015), and it thus seems plausible that damage-induced hormonal changes might have altered the balance of sex-determining hormones in our experiment with *M. annua*. Although *M. annua* has chromosomal sex determination (Russell and Pannell 2015, Veltsos et al. 2018), its sex expression appears to be mediated by the endogenous levels of cytokinin and auxin (Hamdi et al. 1987, Louis et al. 1990, Durand and Durand 1991, Li et al. 2019), and the exogenous application of cytokinins and auxins feminises males and masculinises females, respectively (Hamdi et al. 1987, Durand and Durand 1991). It seems possible that plant responses to herbivory may collaterally affect the cytokinin/auxin ratio resulting in enhanced leakiness, perhaps in an interaction with the phytohormone jasmonate, which has been shown to mediate both anti-herbivore defence (Heil et al. 2001, Thaler et al. 2001, Heil 2008, Kost and Heil 2008, Ballaré 2011) and the sexual development of flowers (Acosta et al. 2009, Wasternack et al. 2013, Cai et al. 2014, Yuan and Zhang 2015, Feng et al. 2020). Furthermore, recent studies on the molecular and genetic mechanisms involved in the sexual expression in persimmon (*Diopsyros kaki*) have shown that the jasmonate pathway and histone methylation/acetylation may play a role in the transition from dioecy to monoecy (Masuda et al. 2020). Although it may seem plausible that enhanced leakiness in damaged *M. annua* plants is pleiotropic response of hormonal signalling, the fact that both males and females became leakier seems to counter such an explanation (because a damage-induced shift in hormone levels or sensitivity ought to shift sex allocation in a particular direction). A more complex relationship between herbivory, phytohormones and sexual expression thus seems necessary, and physiological and/or biochemical analysis of leaky and damaged plants of both sexes would be valuable.

Larger females, but not larger males, were more likely to produce flowers of the opposite sex, or to produce more of them (Table 1). There would seem to be two potential implications for this differential relationship between plant biomass and leakiness in males and females. First, our result reinforces the idea that in *M. annua* male-flower production is costlier than female-flower and fruit production, possibly because of the high investment of nitrogen in pollen (Harris and Pannell 2008, Van Drunen and Dorken 2012, Wright and Dorken 2014). In this context, the fact that we found lower levels of leakiness in smaller individuals only for females is consistent with maleness being the costlier sex in *M, annua*, and with the expectation that larger individuals should allocate more towards the costlier sex because of their greater budget (Delph 1990a, Seger and Eckhart 1996, Klinkhamer et al. 1997, Zhang and Jiang 2002, West 2009). Second, because larger plants haver higher siring success in *M. annua* (Tonnabel et al. 2019), females might benefit from investing in pollen production only when large. In wind-pollinated herbs more generally, large size (in terms of height, which is correlated with biomass) may also directly benefit male function more than female function by promoting the dispersal of pollen from above the plant canopy (Klinkhamer et al. 1997, Friedman and Barrett 2009, Harder and Prusinkiewicz 2013, Tonnabel et al. 2019).

### Implications for plant mating- and sexual-system evolution

Although we remain largely ignorant of how selection might have shaped the interaction between leakiness in sex expression in *M. annua* and its responses to herbivory, there are nevertheless several implications of this interaction for the species’ mating system and potential transitions between sexual systems, which have been frequent in annual lineages of the genus *Mercurialis* (Pannell et al. 2008, Pannell 2018). Most immediately, the induction of higher levels of leakiness by herbivory in our experiment suggests that herbivory in natural populations allow some degree of selfing (and thus mixed mating) in a species that would otherwise be fully outcrossing. We did not estimate the selfing rates in our experimental populations here, but Cossard et al. (2021) found that increased leakiness in females in response to natural selection in experimental populations was associated with up to 22.5% selfing (compared to 0% in dioecious populations). The mating system in *M. annua* is strongly density-dependent, so that even large amounts of pollen produced by females do not lead to high selfing rates in dense populations; indeed most pollen in dense populations competes for outcross siring success (Eppley and Pannell 2007a, Dorken and Pannell 2009, Cossard and Pannell 2019). However, the rate of selfing by leaky females increases rapidly with falling density, so that the production of flowers of the opposite gender by herbivore-damaged plants might have a substantial effect on the mating system in sparse populations.

Our results contribute to a growing picture of the coordinated or interacting nature of mating-system and defence evolution in plants (Carr and Eubanks 2014, Johnson et al. 2015, Lucas-Barbosa 2016). For instance, Johnson et al. (2015) suggested that herbivory could shape the evolution of selfing from outcrossing as a result of herbivore-mediated inbreeding depression, and by affecting pollinator visitation via changes to flowers, potentially leading to pollen limitation. Most previous work has focused on species with combined sexes, and we are unaware of studies on how herbivory might affect reproduction in dioecious species, beyond the observation of (typically) male-biased susceptibility to damage (Ågren et al. 1999, Cornelissen and Stiling 2005, Geber et al. 2012). If herbivory commonly induces leakiness and facultative selfing, as seems to be the case in *M. annua*, then the implications of herbivory for the mating system in dioecious species might differ from that in hermaphroditic species, in which herbivores have been found more typically promote greater outcrossing (Johnson et al. 2015).

Transitions from hermaphroditism to dioecy were long seen as evolutionary dead ends (Heilbuth 2000, Heilbuth et al. 2001, Käfer et al. 2014, Käfer et al. 2017), but recent comparative analysis suggests that reversions to hermaphroditism may have been common (Muyle et al. 2020). In *Mercurialis* in particular, monoecy in polyploid populations is derived from ancestral dioecy, and a study using experimental evolution has demonstrated the role that leakiness in sex expression has likely played in this transition (Cossard et al. 2021). Clearly, hermaphroditism could only ever evolve from dioecy if males or females occasionally expressed both sex functions, either as a result of recombination between sterility loci on a young sex chromosome, thereby regenerating the ancestral hermaphrodite phenotype (Dorken and Barrett 2004, Spigler et al. 2008, Charlesworth 2019, Massonnet et al. 2020), or through leaky sex expression. The expression of leakiness as a result of herbivory would thus represent a potentially interesting case of genetic assimilation, whereby a phenotypically plastic response first exposes a potentially advantageous phenotype to selection (Waddington 1953). If the propensity to respond plastically varies genetically among individuals, as appears to be the case for leakiness in sex expression in *M. annua* (Cossard and Pannell 2019), leakiness induced by herbivory (or mate limitation: Cossard and Pannell 2021) might then quickly become assimilated as established hermaphroditism in response to ongoing natural selection. Further work will be necessary to understand the details of this potential conversion from a plastic to an assimilated state.

## Acknowledgements

We thank Aline Revel for assistance with growing plants.

## Funding

This research was funded by the Swiss National Science Foundation (grant 31003A_163384) and the University of Lausanne.

## Authors contributions

XL and ES collected the data, NV conceived the paper and analysed the data, NV and JRP wrote the paper, with critical inputs from XL and ES.

## Notes

### Competing Interest Statement

The authors have declared no competing interest.

